# Global distribution of earthworm diversity

**DOI:** 10.1101/587394

**Authors:** Helen R P Phillips, Carlos A Guerra, Marie L. C. Bartz, Maria J. I. Briones, George Brown, Olga Ferlian, Konstantin B. Gongalsky, Julia Krebs, Alberto Orgiazzi, Benjamin Schwarz, Elizabeth M. Bach, Joanne Bennett, Ulrich Brose, Thibaud Decaëns, Franciska T. De Vries, Birgitta König-Ries, Michel Loreau, Jérôme Mathieu, Christian Mulder, Wim H. van der Putten, Kelly S. Ramirez, Matthias C. Rillig, David Russell, Michiel Rutgers, Madhav P. Thakur, Diana H. Wall, David Wardle, Data Providers (see bulk upload sheet), Erin Cameron, Nico Eisenhauer

**Affiliations:** German Centre for Integrative Biodiversity Research (iDiv) Halle-Jena-Leipzig, Deutscher Platz 5e, 04103 Leipzig, Germany; Institute of Biology, Leipzig University, Deutscher Platz 5e, 04103 Leipzig, Germany; Universidade Positivo, Rua Prof. Pedro Viriato Parigot de Souza, 5300, Curitiba, PR, Brazil, 81280-330; Departamento de Ecología y Biología Animal, Universidad de Vigo, 36310 Vigo, Spain; Embrapa Forestry, Estrada da Ribeira, km. 111, C.P. 231, Colombo, PR, Brazil, 83411-000; A.N. Severtsov Institute of Ecology and Evolution, Russian Academy of Sciences, Leninsky pr., 33, Moscow, 119071, Russia; M.V. Lomonosov Moscow State University, Leninskie Gory, 1, Moscow, 119991, Russia; European Commission, Joint Research Centre (JRC), Ispra, Italy; University of Freiburg, Tennenbacher Str. 4, 79106 Freiburg, Germany; Global Soil Biodiversity Initiative and School of Global Environmental Sustainability, Colorado State University, Fort Collins, CO 80523 USA; Department of Biology, Colorado State University, Fort Collins, CO, USA; Institute of Biology, Martin Luther University Halle-Wittenberg, Am Kirchtor 1, 06108, Halle (Saale), Germany; Institute of Biodiversity, Friedrich Schiller University Jena, Dornburger-Str. 159, 07743, Jena, Germany; CEFE, UMR 5175, CNRS–Univ Montpellier–Univ Paul–Valéry–EPHE–SupAgro Montpellier–INRA–IRD, Montpellier, France; Institute of Biodiversity and Ecosystem Dynamics, University of Amsterdam, The Netherlands; Institute of Computer Science, Friedrich Schiller University Jena, Ernst-Abbe-Platz 2, 07743 Jena, Germany; Centre for Biodiversity Theory and Modelling, Theoretical and Experimental Ecology Station, CNRS, 09200 Moulis, France; Sorbonne Université, CNRS, UPEC, Paris 7; INRA, IRD, Institut d’Ecologie et des Sciences de l’Environnement de Paris, F-75005, Paris, France; Department of Biological, Geological and Environmental Sciences, University of Catania, Via Androne 81, 95124 Catania, Italy; Department of Terrestrial Ecology, Netherlands Institute of Ecology (NIOO-KNAW), 6700 AB Wageningen, The Netherlands; Netherlands Institute of Ecology, Terrestrial Ecology, 6708PB Wageningen, The Netherlands; Freie Universität Berlin, Institute of Biology, 14195 Berlin, Germany; Senckenberg Museum for Natural History Görlitz, Department of Soil Zoology, 02826 Görlitz, Germany; National Institute for Public Health and the Environment, Bilthoven, The Netherlands; Asian School of the Environment, Nanyang Technological University, Singapore 639798; Faculty of Biological and Environmental Sciences, Post Office Box 65, FI 00014, University of Helsinki, Finland; Department of Environmental Science, Saint Mary’s University, Halifax, Nova Scotia, Canada

## Abstract

Soil organisms provide crucial ecosystem services that support human life. However, little is known about their diversity, distribution, and the threats affecting them. Here, we compiled a global dataset of sampled earthworm communities from over 7000 sites in 56 countries to predict patterns in earthworm diversity, abundance, and biomass. We identify the environmental drivers shaping these patterns. Local species richness and abundance typically peaked at higher latitudes, while biomass peaked in the tropics, patterns opposite to those observed in aboveground organisms. Similar to many aboveground taxa, climate variables were more important in shaping earthworm communities than soil properties or habitat cover. These findings highlight that, while the environmental drivers are similar, conservation strategies to conserve aboveground biodiversity might not be appropriate for earthworm diversity, especially in a changing climate.

**One sentence summary:** Global patterns of earthworm diversity, abundance and biomass are driven by climate but patterns differ from many aboveground taxa.

## Main Text

Soils harbour high biodiversity, and are responsible for a large number of ecosystem functions and services that we rely upon for our well-being (*1*). Despite calls for large-scale biogeographic studies of soil organisms (*2*), global biodiversity patterns remain relatively unknown, with most efforts focused on soil microbes (*3*, *4*), the smallest of the soil organisms. Consequently, the drivers of soil biodiversity, particularly soil fauna, remain unknown at the global scale.

Further, our ecological understanding of global biodiversity patterns (e.g., latitudinal diversity gradients *9*) is largely based on the distribution of aboveground taxa. For many aboveground taxa, variables relating to climate (*10*, *11*) or energy (e.g., primary productivity *12*) are often the most important predictors of diversity across large scales. At large scales, climatic drivers also shape belowground communities (*3*, *13*–*15*), but the response to these drivers in belowground communities may differ from those seen aboveground (*3*, *16*). For example, mean annual temperature correlates positively with aboveground diversity (*17*), but negatively correlates with the diversity of many classes of fungi (*3*), likely due to the optimum temperature of the latter being exceeded (*18*).

Here we analyse global patterns in earthworm diversity, abundance, and biomass (hereafter ‘community metrics’). Earthworms are considered ecosystem engineers (*5*) in many habitats, and increase soil quality (e.g., nutrient availability through decomposition *5*). They also provide a variety of vital ecosystem functions and services (*6*). Whereas most biodiversity-ecosystem functioning studies focus on species richness as a diversity measure (*7*), the provisioning of ecosystem functions by earthworms is likely to vary depending on the abundance, biomass, and ecological group of the earthworm species (*8*) (see Supplementary Materials and Methods). Consequently, understanding global patterns in community metrics for earthworms is critical for predicting how community changes may alter ecosystem functioning.

From small-scale field studies we know that soil properties such as pH and soil carbon influence earthworm diversity (*14*, *19*, *20*). For example, lower pH values constrain the diversity of earthworms by reducing calcium availability (*21*), and soil carbon provides resources that sustain earthworm diversity (*19*). Alongside the many interacting soil properties (*14*), a variety of other drivers can shape earthworm diversity, such as climate and habitat cover (*19*, *22*, *23*). However, to date, no framework has integrated a comprehensive set of environmental drivers of earthworm communities to identify the most important ones at a global scale.

Many soil organisms have shown global diversity patterns that differ from aboveground organisms (*3*, *16*, *24*). Therefore, we anticipate that earthworm community metrics (particularly diversity) will not follow global patterns seen aboveground. This would be consistent with previous studies at smaller scales, which have shown that the species richness of earthworms increases with latitude (*14*, *23*). Because of the relationship between earthworm communities, habitat cover, and soil properties on local scales, we furthermore expect soil properties (e.g., pH and soil organic carbon) to be key environmental drivers of earthworm communities.

Here, we present the first global maps predicting earthworm biodiversity, distilled into three earthworm community metrics: diversity, abundance, and biomass. We collated 181 earthworm diversity datasets from the literature and unpublished field studies (162 and 19, respectively) to create a dataset spanning 56 countries (all continents except Antarctica) and 7048 sites (Fig. 1a). We used these raw data to explore key characteristics of earthworm communities, and determine the environmental drivers that shape earthworm biodiversity. We then used the relationships between earthworm community metrics and environmental drivers (Table S1) to predict local earthworm communities across the globe.

**Fig. 1.**
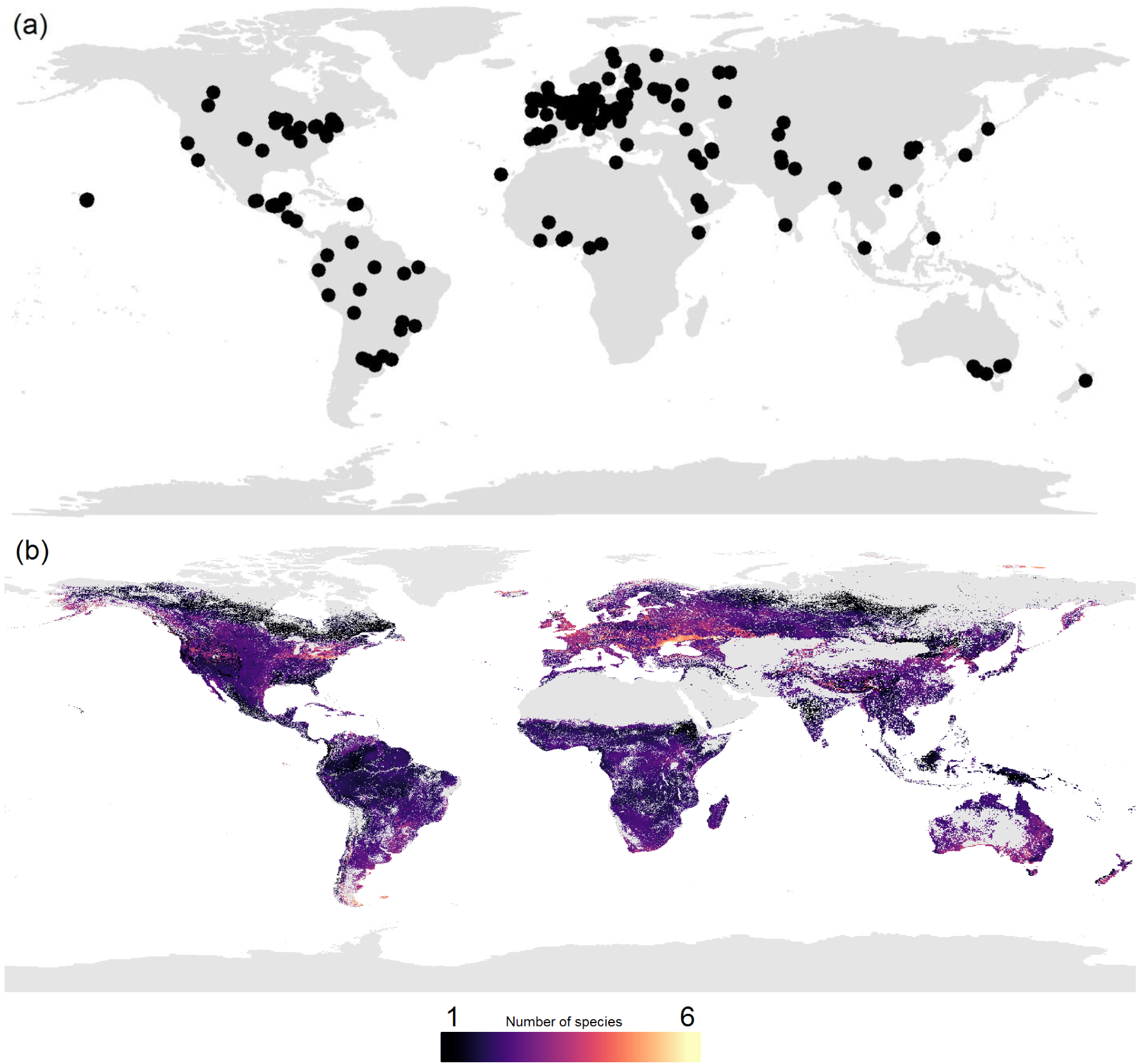
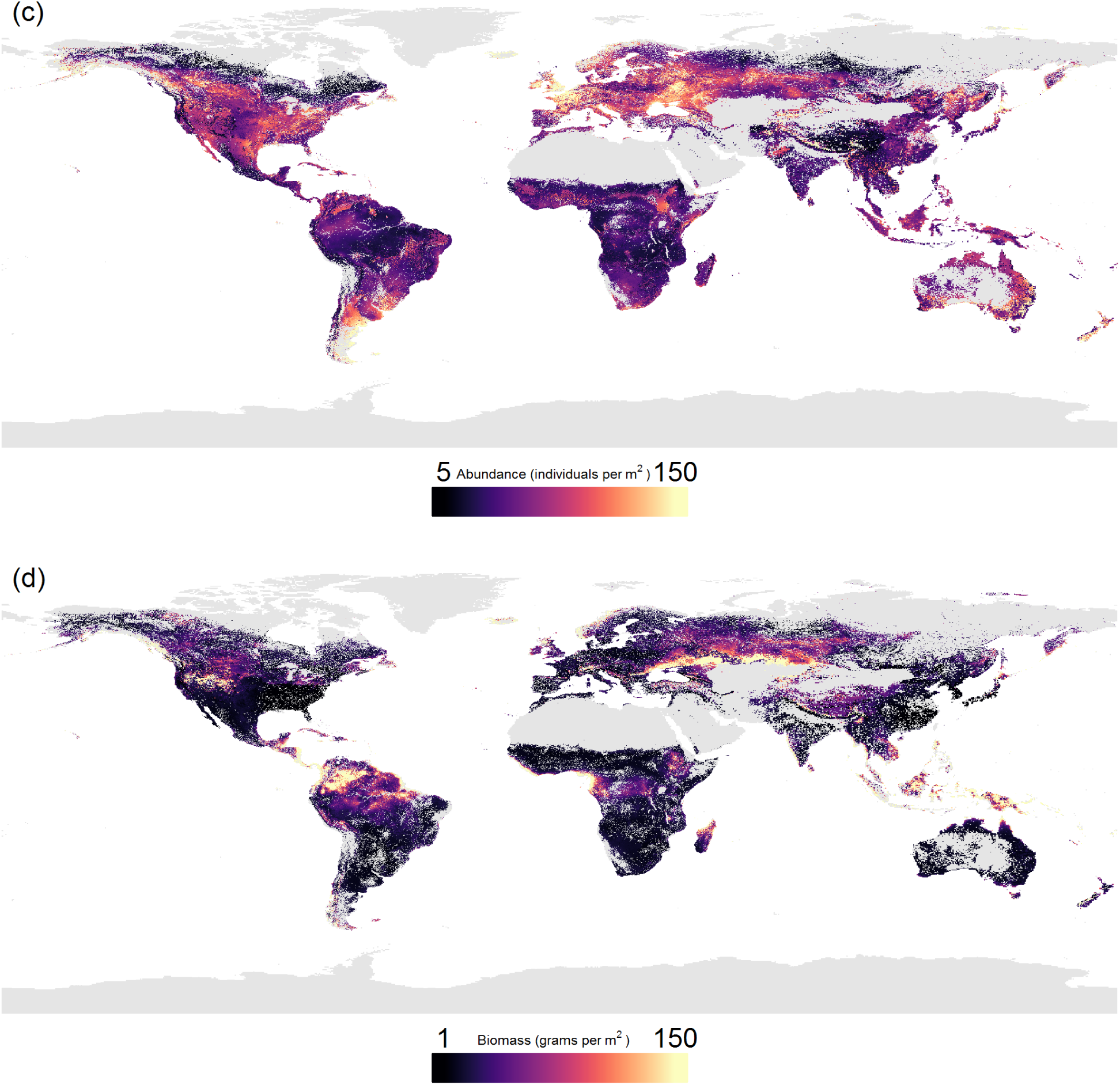
(a) Map of the distribution of data, showing any record that was used in at least one of the three models (species richness, abundance, and biomass). Each black dot represents the centre of a ‘study’ (i.e., a set of data with consistent methodology, see Supplementary Materials and Methods). In total, 229 studies were included (from 181 datasets), which equated to 7048 sites across 56 countries. (b-d): The globally predicted values from the three biodiversity models, species richness (within site, ~1m^2^; panel b), abundance (panel c; individuals per m^2^), and biomass (panel d; grams per m^2^). Areas of high diversity are shown in yellow colours, and areas of low diversity are shown in dark purple colours. Grey areas are habitat cover categories which lacked samples of earthworm communities, thus lack predictions. To prevent outliers skewing the visualization of results, the colour of maps were curtailed at the extreme low and high values. Curtailing was based on where the majority of values laid. Thus, values lower or higher than that number marked on the scale are coloured the same but may represent a large range of values.

Three mixed effects models were constructed, one for each of the three community metrics; species richness (calculated within a site, ~1m^2^), abundance per m^2^, and biomass per m^2^. Each model contained 12 environmental variables as main effects (Table S2), which were grouped into six themes; ‘soil’, ‘precipitation’, ‘temperature’, ‘water retention’, ‘habitat cover’, and ‘elevation’ (see Supplementary Materials and Methods). Within each theme, each model contained interactions between the variables. Following model simplification, all models retained most of the original variables, but some interactions were removed (Table S3).

Predicting based on global environmental data layers, local diversity of earthworms was estimated to range between 1 and 4 species across most of the terrestrial globe (Fig. 1b) (mean: 1.98 species; SD: 0.55). These values are in line with previous suggestions (*20*). The lowest values of species richness occurred across the boreal/subarctic regions, which was expected based on aboveground biodiversity patterns. However, low diversity also occurred in subtropical and tropical areas, such as India and Indonesia, in contrast with commonly observed aboveground patterns, such as the latitudinal gradient in plant diversity. This low earthworm diversity could be due to these regions typically being outside of the optimal temperature range (12-20°C) for earthworms (*25*).

Areas of high local species richness were at mid-latitudes, such as the southern tip of South America, and the southern regions of Australia and New Zealand. Europe (particularly north of the Black Sea) and northeastern USA also had particularly high local species richness. While this pattern seems to contrast with the latitudinal diversity patterns found in many aboveground organisms (*9*, *26*), it is consistent with patterns found in some belowground organisms (ectomycorrhizal fungi *3*, bacteria *15*, *27*), but not all (arbuscular mycorrhizal fungus *13*, oribatid mites *28*). Such mismatches between above- and belowground biodiversity have been predicted (*1*, *24*), but not shown for different soil fauna diversity metrics at the global scale. This work further highlights that it is important that soil organism diversity patterns are examined in concert with those of aboveground taxa if we are to fully understand large-scale patterns of biodiversity and their underlying drivers (*16*, *24*, *29*). Moreover, conservation strategies that are designed for aboveground organisms may not protect earthworms (*24*), despite their importance as ecosystem function providers (*6*) and soil ecosystem engineers (*5*).

The patterns seen here could be a result of past climates, in particular glaciation in the last ice age. Regions in the mid- to high latitudes that were previously glaciated were likely re-colonised by earthworm species with high dispersal capabilities and large geographic ranges (*23*). Thus, mid-latitude communities would have high local diversity but minimal beta-diversity, i.e., low regional diversity, and the opposite would be true in tropical regions. When the number of unique species within each 5-degree latitude band was calculated (i.e., regional richness, Fig. 2a) there was no evidence of a latitudinal diversity gradient once sampling effects had been accounted for (Fig. 2b). Given that regional richness of the tropics was on par with the temperate region, despite low local diversity and relatively low sampling effort (Fig. 2a), endemism of earthworms and beta diversity within the region (*30*) must be considerably higher than within the well-sampled temperate region.

**Fig. 2.**
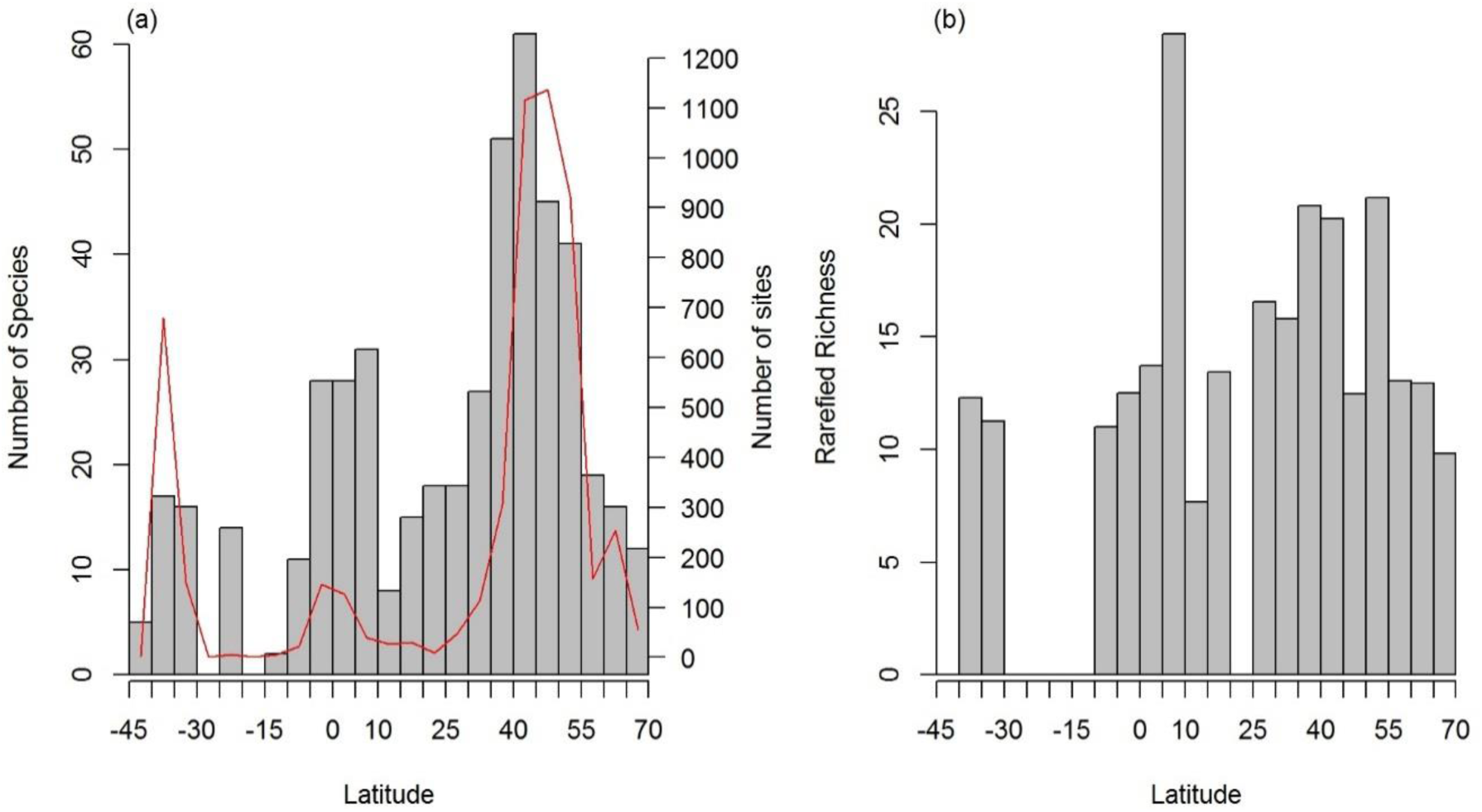
(a) The number of unique species within each 5 degree latitude band (grey bars) and the number of sampled sites within the same latitude band (red line). (b) Sampled-based rarefied species richness within each 5 degree latitude band. Latitude bands with less than 22 sites were not included in the analysis.

Across the globe, the predicted total abundance of the local community of earthworms typically ranged between 5 and 150 individuals per m^2^, in line with other estimates (*31*) (Fig. 1c; mean: 57 individuals per m^2^; SD: 43.59). There was a slight tendency for areas of higher community abundance to be in temperate areas, such as Europe (particularly the UK, France and Ukraine), New Zealand, and part of the Pampas and surrounding region (South America), rather than the tropics. Lower community abundance occurred in many of the tropical and sub-tropical regions, such as Brazil, central Africa, and parts of China. Given the positive relationship between community abundance and ecosystem function (*32*), in regions of lower earthworm abundance there may be implications for provisioning of the ecosystem services performed by these organisms. Further research is needed to disentangle whether these functions are reduced or whether they are carried out by other soil taxa (*1*).

The predicted total biomass of the local earthworm community across the globe typically ranged between 1 g and 150 g per m^2^ (Fig. 1d; mean: 380.86g; SD: 47684.3; median: 18.54, see Supplementary Materials and Methods for discussion in regard to extreme values). The areas of high earthworm biomass were spread across the globe, but concentrated in the tropics (particularly Indonesia, parts of coastal West Africa, Southern Central America, much of Colombia and Western Venezuela), some regions of North America, and the Eurasian Steppe. In some regions, this was almost the inverse of the abundance patterns (Fig. 1c); thus, these results may relate to the fact that earthworms decrease in body size towards the poles (*31*), unlike other animals (*33*). This decrease in earthworm body size might be due to smaller-bodied earthworms with greater dispersal capabilities recolonising northern regions following deglaciation post-ice age (*23*). In northern North America, where there are no native earthworms (*8*), high density and, in some regions, high biomass of earthworms likely reflects the earthworm invasion of these regions. The invasive smaller European earthworm species encounter an enormous unused resource pool, which leads to exceptionally high population sizes (*34*). In contrast, in Brazil, where we had a relatively higher sampling density (Fig. 1a), patterns of abundance and biomass corresponded with the earthworm species that have been documented there. There are a number of giant earthworm species within Brazil (and other countries in the tropics, such as Indonesia, where a similar pattern is shown) (*35*). These giant earthworms normally occur at low densities and low species richness (*35*), causing the high biomass but low abundance.

Overall, the three community metric models performed well in cross-validation (Fig. S2) with relatively high R^2^ values (Table S4 a and c; see Supplementary Material for further details and caveats discussion). But, given the nature of such analyses, models and maps should only be used to explore broad patterns in earthworm communities and not at the fine scale, especially in relation to conservation practices (*12*).

For all three of the community metric models (species richness, abundance, and biomass), climatic variables were the most important drivers (‘precipitation’ theme being the most important for both species richness and total biomass models, and ‘temperature’ theme for the community abundance model; Fig. 3). The importance of climatic variables is consistent with many aboveground taxa (e.g., plants *10*, reptiles/amphibians/mammals *12*) and belowground taxa (bacteria and fungi *3*, *15*, nematodes *27*) when examined at large scales. This suggests that climate-related methods and data, which are typically used by macroecologists for the estimation of aboveground biodiversity, may also be suitable for estimating earthworm communities. However, the strong link between climatic variables and earthworm community metrics is cause for concern, as climate will continue to change due to anthropogenic activities over the coming decades (*36*). Our findings further highlight that changes in temperature and precipitation are likely to influence earthworm diversity (*37*) and their distributions (*14*), with implications for the functions that they provide (*6*). The expansion or shifts in distributions may be particularly problematic in the case of invasive earthworms, such as in areas of North America, where they can considerably change the ecosystem (*8*). However, a change in climate will most likely affect abundance and biomass of the earthworm communities before diversity as shifts in the latter depend upon dispersal capabilities, which are relatively low in earthworms. This underscores the need to study earthworms in terms of multiple community metrics in order to accurately assess responses of communities to climate change.

**Fig. 3:**
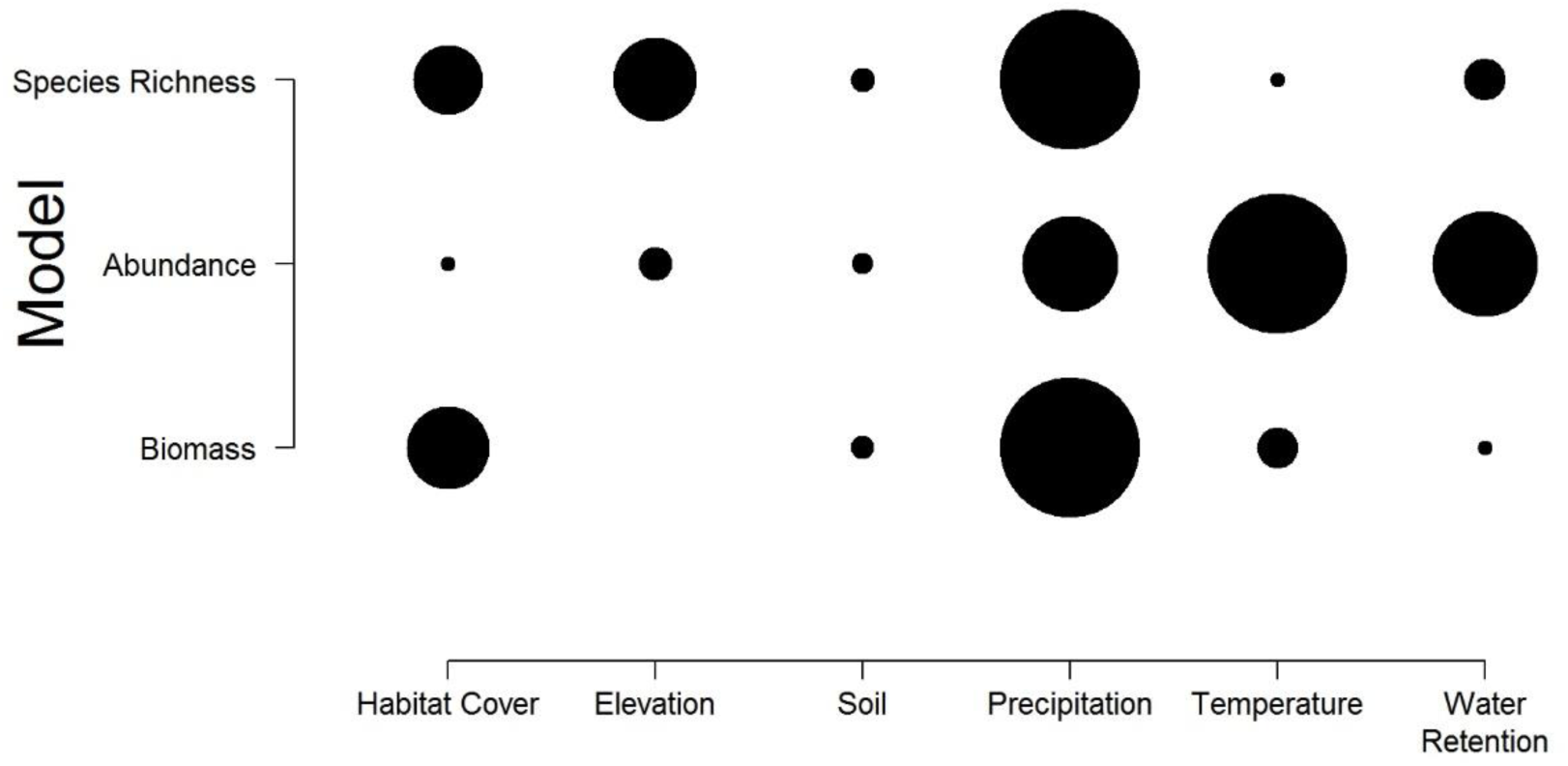
Based on RandomForest models, the importance of the six variable themes from the three biodiversity models. Each row shows the results of each model (top: species richness, middle: abundance, bottom: biomass). Each column represents a theme of variables that was present in the simplified biodiversity model. In the main plot area, the most important variable group has the largest circle. Within each row, the circle size of the other variable themes are scaled in size depending on the relative change in importance. Thus, the circle size should only be compared within a row. Variable theme importance, calculated from the node impurity, was the weighted average of all variables within each theme, following simplification.

We expected that soil properties would be the most important driver of earthworm communities, but this was not the case (Fig. 3). However, soil properties and habitat cover did influence the earthworm community (Fig. S3 a and b). This was especially true in the case of habitat cover, which altered the composition of the three ecological groups (epigeic, endogeics, and anecics, see Supplementary Methods and Materials and Fig. S4) within the earthworm community. Across larger scales, climate influences both habitat cover and soil properties, all of which affect earthworm communities. Being able to account for this indirect effect with appropriate methods and data may alter the perceived importance of soil properties and habitat cover (e.g., with pathway analysis *11* and standardised data). In addition, the importance of drivers could change at different spatial scales, with climate driving patterns at global scales but within climatic regions (or at the local scale) other variables may become more important (*38*). Finally, for soil properties, the mismatch in scale between community metrics and soil properties taken from global layers (for sites where sampled soil properties were missing; see Supplementary Methods and Materials) could also reduce the apparent importance of the theme.

By compiling a global dataset of earthworm communities we show, for the first time, the global distribution of earthworm diversity, abundance, and biomass, and identify key environmental drivers responsible for these patterns. Our findings suggest that climate change might have significant and serious effects on earthworm communities and the functioning of ecosystems. These findings are of particular relevance given the role of earthworms as ecosystem engineers that structure the environment for other soil organisms; thus, any climate change-induced alteration in earthworm communities is likely to have cascading effects on other species in these ecosystems (*8*, *31*). Despite earthworm communities being driven by similar environmental drivers as aboveground communities (*10*, *11*), these relationships result in different patterns of diversity. We highlight the need to integrate belowground organisms into the biodiversity paradigm to fully understand global patterns of biodiversity. This is especially true if the inclusion of soil taxa changes the location of biodiversity hotspots and thus conservation priorities (*24*) or if processes underlying macroecological patterns differ between aboveground and belowground diversity (*29*). Our study creates an avenue for future research: given that climate was the most important predictor of earthworm communities, it is possible for ecologists who have previously focused on modelling aboveground diversity to use similar methods belowground. By modelling both realms, aboveground/belowground comparisons are possible, potentially allowing a clearer view of the biodiversity distribution of whole ecosystems.

## Acknowledgments

We thank Marten Winter and the sDiv team for their help in organizing the sWorm workshops, Peter M. Kotanen, Professor Dr. Jessica G. Davis, S.N. Ramanujam, J.M. Julka, Prof Csaba Csuzdi, P. Bescansa, M. Moriones, C. González, Creighton Litton, Danielle Celentano, Sandriel Sousa, Samuel James, C. Hakseth, C. Mills, Hirohi Takeda, Sandriel Sousa Costa, Kyungsoo Yoo, Sebastien De Danieli, Philippe Choler, Pierre Taberlet, Lauric Cecillon, Erwin Meyer, Dietmar Barkusky, Felix Gerlach, Doris Beutler, Christina Marley, Rhun Fychan, Ruth Sanderson, Mervi Nieminen, Taisto Sirén, Mariana Alem, Carlos Regalsky, Dr. Sandy Smith, Dr. Tara Sackett, Sarah Hamilton, Alexander Rief, Catarina Praxedes, Rosana Sandler, Juliane Palm, Anne Zangerlé, Anne-Kathrin Schneider, Erwin Zehe, Dr. David H. Wise, Dr. Liam Heneghan, Mr. Yoshikazu Kawaguchi, Irene L. López-Sañudo, Almudena Mateos, Pilar Meléndez, Raquel Santos, Marta Yebra, Tamara Vsevolodova-Perel, Maxim Bobrovsky, Natalya Ivanova, and Eufemio Rasco Jr..

## Funding

This work was developed during and following two ‘sWorm’ workshops. HRPP and the sWorm workshops were supported by the sDiv (Synthesis Centre of the German Centre for Integrative Biodiversity Research (iDiv) Halle-Jena-Leipzig (DFG FZT 118)). ML was supported by the TULIP Laboratory of Excellence (ANR-10-LABX-41). KSR and WvdP were supported by (ERC-ADV grant 323020 to WvdP). In addition, data collection was funded by: Russian Foundation for Basic Research (19-05-00245), Tarbiat Modares University, Fonds de Recherche du Québec – Nature et technologies, Aurora Organic Dairy, UGC(NERO) (F. 1-6/Acctt./NERO/2007-08/1485), Natural Sciences and Engineering Research Council (RGPIN-2017-05391), Slovak Research and Development Agency (APVV-0098-12), Science for Global Development through Wageningen University, Norman Borlaug LEAP Programme and International Atomic Energy Agency (IAEA), São Paulo Research Foundation - FAPESP (12/22510-8), Oklahoma Agricultural Experiment Station, INIA - Spanish Agency (SUM 2006-00012-00-0), Royal Canadian Geographical Society, Environmental Protection Agency (Ireland) (2005-S-LS-8), University of Hawai’i at Manoa (HAW01127H; HAW01123M), CAPES, European Union FP7 (FunDivEurope, 265171), U.S. Department of the Navy, Commander Pacific Fleet (W9126G-13-2-0047), Science and Engineering Research Board (SB/SO/AS-030/2013), Strategic Environmental Research and Development Program (SERDP) of the U.S. Department of Defense (RC-1542), Maranhão State Research Foundation (FAPEMA), Coordination for the Improvement of Higher Education Personnel (CAPES), Ministry of Education, Youth and Sports of the Czech Republic (LTT17033), Colorado Wheat Research Foundation, Zone Atelier Alpes, French National Research Agency (ANR-11-BSV7-020-01, ANR-09-STRA-02-01), Austrian Science Fund (P16027, T441), Rentenbank, Welsh Government and the European Agricultural Fund for Rural Development (Project Ref. A AAB 62 03 qA731606), SÉPAQ, Ministry of Agriculture and Forestry of Finland, McKnight Foundation, Science Foundation Ireland (EEB0061), University of Toronto (Faculty of Forestry), National Science and Engineering Research Council of Canada, Haliburton Forest & Wildlife Reserve, NKU College of Arts & Sciences Grant, Österreichische Forschungsförderungsgesellschaft (837393 and 837426), Mountain Agriculture Research Unit of the University of Innsbruck, Higher Education Commission of Pakistan, Kerala Forest Research Institute, Peechi, Kerala, UNEP/GEF/TSBF-CIAT Project on Conservation and Sustainable Management of Belowground Biodiversity, Ministry of Agriculture and Forestry of Finland, Complutense University of Madrid / European Union FP7 project BioBio (FPU UCM 613520), GRDC, AWI, LWRRDC, DRDC, CONICET (National Scientific and Technical Research Council) and FONCyT (National Agency of Scientific and Technological Promotion) (PICT, PAE, PIP), Universidad Nacional de Luján y FONCyT (PICT 2293 (2006)), Fonds de recherche sur la nature et les technologies du Québec (131894), Deutsche Forschungsgemeinschaft (SCHR1000/3-1, SCHR1000/6-1, 6-2 (FOR 1598), WO 670/7-1, WO 670/7-2, & SCHA 1719 / 1-2), CONACYT (FONDOS MIXTOS TABASCO/PROYECTO11316), NSF, Institute for Environmental Science and Policy at the University of Illinois at Chicago, Dean’s Scholar Program at UIC, Garden Club of America, the Nature Conservancy, The College of Liberal Arts and Sciences at Depaul University, and Elmore Hadley Award for Research in Ecology and Evolution from the UIC Dept. of Biological Sciences (DGE?0549245, DEB?BE?0909452, NSF1241932), Spanish CICYT (REN2000-0783/GLO; REN2003-05553/GLO; REN2003-03989/GLO), Grant-in-Aid from the Graduate School of Environment and Information Science of Yokohama National University, Grant-in-Aid for the Global COE program E03 for Global Eco-Risk Management from Asian Viewpoints, YNU International Environmental Leaders Program in Sustainable Living with Environmental Risk (SLER) funded by Promoting Science and Technology System Reform of the Promotion of Science and Technology, Japan (MEXT KAKENHI Grant Number 25220104, JSPS KAKENHI Grant Number 25281053), ADEME (0775C0035), Ministry of Science, Innovation and Universities of Spain (CGL2017-86926-P), Syngenta Philippines, UPSTREAM, Japan Society for the Promotion of Science (JSPS KAKENHI Grant Number 17TK0074), LTSER (Val Mazia/Matschertal).

Author contributions

Competing interests:

Data and materials availability:

## Supplementary Materials

Materials and methods

Table S1 – S4

Figures S1 - S4

