## Supplementary Material and Methods for "Global distribution of earthworm diversity" (from another version of this preprint: version 1, 2019-03-24)

**Literature Search**

Web of Science was searched on 18th December 2016, using the following search term:

*((Earthworm* OR Oligochaeta OR Megadril* OR Haplotaxida OR Annelid* OR Lumbric* OR Clitellat* OR Acanthodrili* OR Ailoscoleci* OR Almid* OR Benhamiin* OR riodrilid* OR Diplocard* OR Enchytraeid* OR Eudrilid* OR Exxid* OR Glossoscolecid* OR Haplotaxid* OR Hormogastrid* OR Kynotid* OR Lutodrilid* OR Megascolecid* OR Microchaetid* OR Moniligastrid* OR Ocnerodrilid* OR Octochaet* OR Sparganophilid* OR Tumakid* ) AND (Diversity OR "Species richness" OR "OTU" OR Abundance OR individual* OR Density OR "tax* richness" OR "Number" OR Richness OR Biomass))*

This search returned 7783 papers. All titles and abstracts of papers post-2000 were screened (6140 papers), and were excluded if they did not make reference to data suitable for the analysis (suitability discussed below). As it was most likely that raw data would need to be requested, papers published before 2000 were not screened, as it was unlikely that available author contact details were up-to-date. We note however that earlier publications may be useful for future research, e.g., focusing on long-term monitoring. After this initial screening, PDFs of all remaining papers (n = 986) were manually screened to determine whether data were suitable.

In order to be suitable for the analysis, papers had to present (or make reference to) the following information and data:

1. Sampled earthworm communities using an appropriate earthworm extraction methodology, such as hand-sorting of a sufficient soil volume e.g. (Anderson and Ingram 1993) or chemical expulsion from a quadrat e.g., (ISO 2018) at two or more sites. At a

minimum, total fresh biomass and/or total abundance of the earthworms at each site had to be measured. Ideally, there was data on identification of all individuals to species level, with the abundance/biomass data of each species;

2. Available geographic coordinates for all sampled sites, or maps that could be georeferenced;
3. Measurements of at least one soil property at each site (see below);
4. Information on the habitat cover and/or land use;
5. Differences in land use/habitat cover or soil properties (see below for information on the land use/habitat cover and soil properties) across the sites.

Where possible, all suitable data were taken from the 477 papers that were identified as containing suitable data. Data were extracted from figures where necessary (using IMAGEJ, Schindelin et al. 2012). If data was not provided in the text or the supplementary materials, authors were contacted to request the raw data from each site. As some datasets remain unpublished, or are yet to be published, individual earthworm researchers were also contacted to enquire as to whether they had suitable data. Including unpublished data helps to reduce publication bias (Koricheva et al. 2013).

**Data collation**

The data taken or requested from one publication or an unpublished field campaign was considered a ‘dataset’. If a dataset contained data sampled using different methodologies, we split it into different ‘studies’ based on the methodology, as measured diversity of earthworms is highly dependent on the methods used; (Bartlett et al. 2010). For datasets where sites were repeatedly sampled over time, both within years and across years, we used only the first and the last sampling campaign, and these were split into two studies. The modelling approach used (mixed-effects models, with random effects accounting for different studies) dealt with non-independence of such datasets (Crawley 2012).

*Site level information*

For each dataset, we collated the following information into a standardised data template: geographic coordinates for each of the sampled sites, start and end dates of sampling (month and year), and the sampling method used. For each dataset, we requested at least one soil property (pH, cation exchange capacity (CEC) or base saturation, organic carbon, soil organic matter, C/N ratio, soil texture, soil type, soil moisture) for each site. If soil properties were given for different soil depths, then we calculated a weighted average (maximum soil depth = 1 m, but typically collected down to 30 cm). Using information within the published articles, and additional information provided by the data collectors, the habitat cover at each site was classified into categories based on the ESA CCI-LC 300m map (http://maps.elie.ucl.ac.be/CCI/viewer/index.php; Supplementary Table 1).

*Recorded community metrics*

For each dataset, the following site-level community metrics were calculated where possible: total abundance of earthworms at the site, total fresh biomass of earthworms at the site and number of species at the site. Using the area sampled at the site, both abundance and biomass were transformed to individuals per $m^2$ and grams per $m^2$, respectively, if they were not already given in that unit. Species richness of the site was calculated from available species lists, if not already provided. Two issues arose when calculating species richness of earthworms. Firstly, many specimens were not identified to species level. Where data collectors identified a specimen as a unique morphospecies (species delineation based solely on morphological characteristics), they were included in the species richness estimate as an additional species. Where unidentified species were grouped as ‘unknown’, these were excluded. Secondly, typically only adult specimens of many earthworm species can be identified to species level (Bartlett et al. 2010), so juveniles were excluded from the calculation. Therefore, a more appropriate term would be ‘number of identified adult (morpho-) species’, but for brevity this will be referred to as ‘species richness’.

*Species identity*

For datasets where the earthworms had been identified to species levels, all species names were checked for spelling errors and synonyms. Scientific names were standardised using expert opinion (MJB, GB, MB, PL) and DriloBASE (http://drilobase.org/drilobase). Following standardisation, earthworm species were categorised into the three main ecological groups: epigeics, endogeics, and anecics (Bouché 1977), plus a fourth minor group, epi-endogeic (species which exhibit traits of both epigeics and endogeics). Epigeic species are typically found in the upper layers of the soil and litter, and, amongst other roles, are important in the first stages of decomposition through burial of the litter layer (Brown 1995, Edwards 2004, Seeber et al. 2008). Endogeic species live in the upper mineral soil layers, creating horizontal burrows (Bouché 1977). One function they have been shown provide is a decrease the density of root-pathogenic nematodes (Blouin et al. 2005, Boyer et al. 2013), reducing nematode populations and disease incidence, thereby increasing crop yields (Brown et al. 2004, Loranger-Merciris et al. 2012). Anecic species mix the litter and mineral soil via surface cast production (Bouché 1977, Brown 1995). In addition, the vertical burrows created by anecic species increase water infiltration into deeper soil layers, increasing water holding capacity (Joschko et al. 1989, Bouché and Al-Addan 1997), and regulating water availability.

**Data extraction and harmonization across global layers**

In order to predict earthworm communities across the globe, we required harmonised sets of spatially distributed variables. We collected 15 globally distributed layers that are described as predictors of earthworm distribution (Supplementary Table 2). For the SoilGrids data (Hengl et al. 2017; https://soilgrids.org), which provides soil properties for different layers within the soil profile, we calculated the weighted average for the values of the top four layers (corresponding to the top 30 cm of the soil profile, which matches with most of the earthworm sampling techniques). For sites missing sampled soil properties, the soil properties associated with the 1km pixel corresponding to the site's geographical coordinates were used in the analyses.

Where possible, the land cover global layer (ESA CCI-LC 300m; https://www.esa-landcover-cci.org/) was re-categorised to amalgamate similar habitat cover categories matching the ones collected within the dataset (see Supplementary Table 1). Where not possible, the categories were ignored (i.e., classified as NA) during later steps, as estimates could not be produced for unknown habitat cover categories.

No climate variables were taken from the papers or raw data provided. Instead, five global climate layers (climatologies) obtained from the CHELSA climate dataset (Karger et al. 2017) were used (annual mean temperature, temperature seasonality, temperature annual range, annual precipitation, and precipitation seasonality) and, from other sources, the number of months of snow cover (Hall and Riggs 2015), and the aridity index and potential evapotranspiration (PET; Zomer et al. 2007, 2008). The within-year standard deviation of PET (PETSD) was calculated as well. Finally, a globally distributed layer of elevation (Danielson and Gesch 2011) was also included in the analysis. For all of these layers, the value within the 1km pixel that matched the site's coordinates was used in the analyses.

For an initial harmonisation across all global layers, it was necessary to aggregate or disaggregate - when appropriate - the spatial resolution of the different layers to match a one kilometer square grid. A nearest neighbour disaggregation algorithm was applied without changing the pixel values, but changing the pixel resolution using the one kilometer square resolution from SoilGrids as a reference.

Following the spatial harmonization, the global layers were matched with the collated dataset based on the geographic coordinates of the sampled sites. In the case of the climate layers, all variables were appended to the dataset. Soil variables were only appended if the sites were missing sampled measures (most sites contained pH values, 62.7%, but only 7.3% contained CEC, for example).

To help prevent extrapolation, all global layers were truncated to values represented by each subset of data, i.e., the minimum and maximum values used in each of the three community metric models. The exception was the number of months of snow cover, which was truncated at four months, thus any sites or areas of the globe with a greater number of months than four were modelled and predicted (respectively) as four months. This ensured an even spread across the range of values (many sites were within 0-4 months, only 9% of sites were greater than four).

**Mixed effects modelling**

*Earthworm species richness, abundance and biomass models*

Three mixed effects models were constructed, using lme4 (Bates et al. 2015), one for each of the site-level community metrics: species richness, total abundance (individuals per $m^2$) and total biomass (grams per $m^2$). Prior to modelling, the full dataset was split into three subsets, based on the response variables (i.e., a dataset containing all sites with a species richness value). Within each dataset, we tested for multicollinearity between the elevation, climate and soil variables using Variance Inflation Factors (VIFs) and removing the variable with highest VIF in turn until all remaining variables were below the predetermined threshold of 3 (Zuur et al. 2010).

Abundance and biomass were log transformed (ln + 1) prior to modelling and were then modelled using a Gaussian error structure. Species richness was not log transformed, but instead modelled with a Poisson error structure. All models had random effects that accounted for variation between each of the different studies, using an intercept only structure. Fixed effects included habitat cover, elevation, soil properties and climate variables. All continuous variables (i.e., elevation, all soil variables and most climate variables) were centred and scaled (variables were centred on the mean value and divided by the standard deviation) to aid model fitting and interpretability. Number of months of snow cover was modelled as a categorical variable (and therefore not centred and scaled) to allow for a non-linear relationship. As it is expected that earthworm diversity will peak with some snow cover, due to increased precipitation, and soil protection during freezing months (Eisenhauer et al. 2014), but be restricted by prolonged snow cover (Nieminen et al. 2011). This also improved the modelling process, as sites were skewed towards the lower number of months, with not enough data at the higher latitudes to fit a non-linear regression.

For each of the three models, the structure of the fixed effects in the maximal model was the same. Habitat cover and elevation were included as additive effects with no interactions. The other variables were grouped into four themes; ‘soil’, ‘precipitation’, ‘temperature’ and ‘water retention’ (Supplementary Table 3). For example, all precipitation variables that remained (i.e., were not removed due to their VIF score) were grouped together. Within the soil and two climate groups, all two-way interactions were considered. The water retention group contained specific two-way interactions between soil structure variables (clay and silt percentage) and climate variables relating to water availability that were present in the two climate themes (annual precipitation, precipitation seasonality, PET, PETSD, and aridity). These specific interactions were to account for soil moisture and how quickly moisture might leave the soil.

Each maximal model was then simplified using akaike information criterion (AIC) values. All interactions were tested first, and removed if AIC values were reduced compared to the more complicated model. Any main effects that were not involved with interactions were tested, and removed if AIC values were reduced (Zuur et al. 2009, Crawley 2012) (Supplementary Table 3).

*Ecological group responses*

The same modelling approach was used to investigate changes in earthworm ecological groups across the different habitat types. For each site, the biomass of the three main ecological groups (epigeic, endogeic, and anecic) and one minor ecological group (epi-endogeic) were calculated, based on the category assigned following species name standardisation. Mixed effects models were constructed as detailed above, with the exception that habitat cover interacted with the ecological group, with ecological group biomass at the site as a response variable. The model was simplified following details given above.

The biomass of each ecological group in each habitat cover was then predicted, using the ‘predict’ function in ‘lme4’ (when all other variables were at zero, i.e., the mean). The predicted values for the three main ecological groups, (epigeic, endogeic and anecic, which had sufficient underlying data) were plotted using the ‘triangle.plot’ function in ‘ade4’ (Dray and Dufour 2007). The predicted total biomass, i.e., the total of the predicted biomass of the three main ecological groups, was used to determine size of the points within the triangle plot.

**Creating maps of earthworm communities**

The global patterns of earthworm communities (species richness, abundance, and biomass) were predicted using each of the three models. The values from the relevant global layers (i.e., those corresponding to the variables that remained in each model following simplification) were used in the 'predict' function in the 'lme4' package, being predicted based on the coefficients of the final models.

A global layer of predicted values was then presented as maps of local communities of earthworms. Although all global layers had been capped at values represented in the underlying dataset, extrapolation still occurred during the prediction (there were instances where grid cells in multiple layers were at the extreme values, and such combinations were not represented in the underlying data, most evident in the predictions of earthworm biomass, see 'Interpreting the model validation').

**Variable Importance**

In order to determine which themes (soil, habitat cover, precipitation, temperature, water retention) were the most influential in driving earthworms communities, Variable Importance was performed using RandomForest models (Breiman 2001, Liaw and Wiener 2002).

For each of the three community metrics, random forest models were constructed (Liaw and Wiener 2002) with all the variables that were present in the final (i.e. simplified) model. Random forest models use multiple regression trees to classify data (Liaw and Wiener 2002). This method was chosen as these models can handle non-linear data, whilst interactions are not specified but can be learnt from the data (Grömping 2009). Random forest models are an ensemble of individual regression (or classification) trees (Breiman 2001, Liaw and Wiener 2002). Each tree is created using only a subset of the available data, i.e., "out-of-bag" regression, and the process is repeated until the 'forest' is complete (ntree default = 500 trees).

At each node in the tree, the subset of response variables is split using the best predictor variable. Unlike regression trees, where at each node the best predictor is used from all available predictor variables, random forest models use only a random sample of the predictor variables (“Mtry”) to determine the best predictor to split the response variable at each node (Breiman 2001, Liaw and Wiener 2002). The default Mtry value was used (number of predictors divided by 3), so in our case of 10 to 12 predictor variables Mtry = 3 (biomass model) and 4 (species richness and abundance models). The “out-of-bag” data is then predicted using the average prediction of all trees (Liaw and Wiener 2002).

In addition, random forest models can be used to assess the importance of individual variables (Breiman 2001). One such measure is the mean decrease in node impurity calculated from the decrease in the residual sum of squares for the variable that was used at the node. The average decrease for each variable is averaged across all the trees to create the node impurity (Liaw and Wiener 2002). An alternative importance variable is the mean decrease in accuracy. For each tree, when the “out-of-bag” data is being predicted, a single predictor variable is permuted, and the increase in prediction error calculated (Liaw and Wiener 2002). This mean decrease in accuracy is often considered the best of those available (Strobl et al. 2007), but results between the decrease in node impurity and mean decrease in accuracy often correlate well (Menze et al. 2009).

For each of the three random forest models, the mean decrease in node impurity and mean decrease in accuracy was calculated (using the ‘importance’ function in ‘randomForest’) for each predictor variable in the random forest. In order to determine which theme of variables (habitat cover, elevation, soil, temperature, climate, and water retention) was most important in driving patterns in earthworm communitiess, the mean decrease in node impurity was averaged for all variables within each theme and weighted by the number of times each variable was used in the random forest compared to the other variables in the same theme.

**Model validation and sensitivity analysis**

A number of additional analyses were performed to determine the robustness of the models and the ability to predict new values. Firstly, the influence of combining both measured soil properties and soil properties from SoilGrids was tested. For the three response variables (species richness, abundance, and biomass) models were created that only included data from SoilGrids. The same modelling process was used as described above (using VIFs to determine starting variables, then simplification). Once the final model had been identified for each of the three community metrics, 10-fold cross-validation was performed.

Cross-validation was performed by randomly splitting the dataset underlying each of the three models into 10 nearly-equal size groups. Using the model structures produced following simplification, the model was built using 9 of the groups of data. The 10th group of data was predicted from the re-built model. The predicted data was plotted against the observed data. This process was repeated until all 10 groups of data had been predicted. This process was done for the models that contained only SoilGrids data, and the main models (that used a mixture of soil property data).

For the cross-validated models, the mean squared error (MSE) was calculated from the results of the cross-validation. MSE measures the ability of the model to predict new data, and the result are easily interpretable as they are on the same scale as the original data. MSE was calculated for the total of all models, but as the models may be better at predicting certain ranges of values, MSE was also calculated for the tertiles of the observed data (i.e., the ability of the model to predict the low, medium and high values of earthworm communities).

Secondly, the $R^2$ values of all models (the main models, and the models with only SoilGrids data) were calculated using the MuMIn package (Barton 2018). The $R^2$ values describe the fit of the model to the data.

**Latitudinal diversity gradients**

In order to ascertain whether there is a species diversity gradient with latitude, the site-level diversity data underlying the species richness model (i.e., contained sites with species level or morphospecies identification) was used. The sites were split into five degree latitude bands from the equator (to maximise number of bands whilst minimising the number of bands with no samples), and the number of unique species within each band was calculated. Within each band, it was assumed that each uniquely named morphospecies was different from any of the named species (number of morphospecies across bands, min = 0, max = 14, mean = 2.6). Some of the sites also contained genus-level only identification. When this was the case, a genus was included as an additional species if the genus was unique within the band (i.e., no named species belonged to that genus). The number of sites within each latitude band was also calculated.

As the number of sites, a measure of sampling intensity, differed a large amount across each band (minimum = 3, maximum = 1136), sample-based rarefied richness was calculated (Gotelli and Colwell 2010). Within each band, sample-based rarefied richness was calculated as the mean number of species (including morphospecies, and unique genus', as described above) across 1000 random samples (without replacement) of *n* sites. Typically, *n* would be the minimum number of sites within a latitude band (in the case of this dataset, *n*=3). However, this was deemed too low, and so *n*=22 was used (the fourth lowest sampling intensity within a band. Other n's are produced, presented in Supplementary Figure 1, but did not change the conclusions). Where the number of sites within a band was less than *n*, the rarefied richness of the band was not calculated. By removing the effect of sampling intensity, any latitudinal diversity gradient could be attributed to other factors, such as land area or climate.

All statistics, data manipulation and processing of global data layers was implemented in R (version 3.3.1; R Core Team 2016). All code is available [DOI].

**Interpreting the model validation**

The results of the biomass model highlighted an issue with the modelling technique used. All of the global data layers were cut at values represented by the underlying datasets. However, during the prediction, it was often the case that multiple data layers were at the extreme ends of the possible range of values. This led to, especially in the case of the biomass model, and to some extent the abundance model, unrealistic values being predicted. This issue would only be fixed with additional data, but would not affect the visual maps produced.

Overall, the models had reasonably good fit to the data, assessed using the $R^2$ values (Supplementary Table 4c). However, the predictive power of the models was variable. With all models, the total MSE (Supplementary Table 4a) increased mainly due to the ill-fitting of the sites with higher values. It is unclear why high values cannot be fitted well with the models; however, it is highly likely that increasing the number of sites would help either identify the issue or improve model fit.

For the majority of the datasets (182 out of 229 studies), the models contained the measured soil properties for some of the variables. However, where this was missing, we used the SoilGrids data. Models which contained only SoilGrids data had a better fit to the data (Supplementary Table 4c) and were typically better at predicting during cross-validation (lower

MSE values; Supplementary Table 4b). However, in most instances, the change in MSE was negligible between the different types of models (Supplementary Table 4 a versus b). However, there are many reasons why using a mixture of the measured variables and the SoilGrids variables is the best option, rather than just using SoilGrids data. Firstly, modelled global estimates of the soil properties may not accurately depict site-level conditions (Sanchez et al. 2009), which could result in the variables appearing less important than they would be if they matched the measured communities. Secondly, some of the coordinates within a study were identical which would result in identical SoilGrids data (for these datasets, often small-scale field experiments, the measured soil properties variables were not identical). Using only SoilGrids data would reduce the gradient of soil properties within each study, reducing the number of available gradient comparisons across all datasets. And given that a number of studies (106 out of 229 studies) had identical climate variables across all sites, having variety in all other variables prevented this being in an issue within the modelling framework. We call on soil ecologists to collect data on soil properties when they measure diversity of soil taxa, as this permits more robust modelling at both the small scale, and across larger scales.

Regardless of whether the model contained measured soil properties or only SoilGrids data, the models were consistently worse at predicting when observed values was high. This is likely due to the small number of studies where sampled values were high. Only 5 studies had more than 10 species of earthworms in at least one site, and only 6 studies had more than 300 grams per $m^2$ of earthworm biomass in at least one site. There were a greater number of studies that contained high abundance of earthworms, with 34 studies having at least one site that contained more than 600 individuals per $m^2$. Increasing the number of studies and sites would help identify whether this, or another cause, is responsible. Ideally, this would improve the predictive power of the models. It is hoped that efforts will continue to collate earthworm diversity data from across the globe.

### Supplementary Tables

**Supplementary Table 1**: The re-categorisation of the ESA habitat cover variable. Habitat cover at a sampled site was classified based on the ‘Reclassified habitat cover’ column. As not all categories of habitat would be available in the data (i.e., due to too detailed categories, or in habitats typically devoid of sampling), some of the categories of the original habitat cover variable (left-hand column) were reclassified (right-hand column). Usually, this means that categories were grouped together (i.e., to reduce the categories based on ‘openness’).

| **Original habitat cover variable** | **Reclassified habitat cover** |
|---|---|
| No Data | NA |
| Cropland, rainfed | Production - Herbaceous |
| Cropland - herbaceous cover | Production - Herbaceous |
| Cropland - Tree or shrub cover | Production - Plantation |
| Cropland, irrigated or post-flooding | NA |
| Mosaic cropland (>50%) / natural vegetation (tree, shrub, herbaceous cover) (<50%) | Cropland/Other vegetation mosaic |
| Mosaic natural vegetation (tree, shrub, herbaceous cover) (>50%) / cropland (<50%) | Cropland/Other vegetation mosaic |
| Tree cover, broadleaved, evergreen, closed to open (>15%) | Broadleaf evergreen forest |
| Tree cover, broadleaved, deciduous, closed (>40%) | Broadleaf deciduous forest |
| Tree cover, broadleaved, deciduous, open (15-40%) | Broadleaf deciduous forest |
| Tree cover, needleleaved, evergreen, closed (>40%) | Needleleaf evergreen forest |
| Tree cover, needleleaved, evergreen, open (15-40%) | Needleleaf evergreen forest |
| Tree cover, needleleaved, deciduous, closed (>40%) | Needleleaf deciduous forest |
| Tree cover, needleleaved, deciduous, open (15-40%) | Needleleaf deciduous forest |
| Tree cover, mixed leaf type (broadleaved and needleleaved) | Mixed forest |
| Mosaic tree and shrub (>50%) / herbaceous cover (<50%) | Tree open |
| Mosaic herbaceous cover (>50%) / tree and shrub (<50%) | Herbaceous with spare tree/shrub |
| Shrubland | Shrub |

| | |
|---|---|
| Grassland | Herbaceous |
| Lichens and mosses | NA |
| Sparse vegetation (tree, shrub, herbaceous cover) (<15%) | Sparse vegetation |
| Tree cover, flooded, fresh or brakish water | NA |
| Tree cover, flooded, saline water | NA |
| Shrub or herbaceous cover, flooded, fresh/saline/brakish water | NA |
| Urban areas | Urban |
| Bare areas - consolidated | Bare area (consolidated) |
| Bare areas - unconsolidated | Bare area (unconsolidated) |
| Water bodies | Water bodies |
| Permanent snow and ice | NA |

**Supplementary Table 2**: Information for each of the 15 global layers detailed in the methods.

| | ***Variable*** | ***Source*** | ***Original Spatial Resolution*** |
|---|---|---|---|
| | Habitat Cover | ESA CCI-LC | 300m |
| | Elevation | (Danielson and Gesch 2011) | 1km |
| Soil Parameters | pH (H20) | SoilGrids (Hengl et al. 2017) | 1km |
| | Organic carbon | | 1km |
| | Soil clay content | | 1km |
| | Soil silt content | | 1km |
| | CEC | | 1km |
| Temperature | Annual Mean Temperature | CHELSA (Karger et al. 2017) | 1km |
| | Temp. seasonality | | 1km |
| | Temp. annual range | | 1km |
| | PET | (Zomer et al. 2007, 2008) | 1km |
| | PETSD | | 1km |
| Precipitation | Annual precipitation | CHELSA (Karger et al. 2017) | 1km |
| | Precip. seasonality | | 1km |
| | Number of Months with Snow | (Hall and Riggs 2015) | 1km |
| | Aridity Index | (Zomer et al. 2007, 2008) | 1km |

**Supplementary Table 3:** Results following model simplification of the three community metric models. Highlighted cells show which interactions remained following simplification for the species richness (pink), abundance (blue) and biomass (green) models. Some variables were only present in the simplified model as main effects, indicated in the column 'Main Effect Only'. Also noted is the variable theme in which the variable was grouped. Variables that interacted within the 'water retention' theme are not shown explicitly, but can be deduced based on interactions between a climate variable and soil property variable. Abbreviations: CEC = Cation exchange capacity, Temp. = Temperature, Precip. = Precipitation, PET = Potential evapotranspiration, PETSD = within year standard deviation of PET

| ***Global Data Layers*** | | Main Effect Only | pH ($H_2O$) | Organic carbon | Clay content | Silt content | CEC | Annual Mean Temperature | Temp. seasonality | Temp. annual range | PET | PETSD | Annual precipitation | Precip. seasonality | Number of Months with Snow |
|---|---|---|---|---|---|---|---|---|---|---|---|---|---|---|---|
| | Habitat Cover | pink, blue, green | | | | | | | | | | | | | |
| | Elevation | pink, blue | | | | | | | | | | | | | |
| *Soil* | pH ($H_2O$) | | | | | | | | | | | | | | |
| | Organic carbon | | pink, blue, green | | | | | | | | | | | | |
| | Clay content | | green | blue | | | | | | | | | | | |
| | Silt content | | blue, green | pink | | | | | | | | | | | |
| | CEC | | pink, blue, green | pink, blue, green | pink, blue, green | | | | | | | | | | |
| *Temperature* | Annual Mean Temperature | | | | | | | | | | | | | | |
| | Temp. seasonality | | | | | | | | | | | | | | |
| | Temp. annual range | | | | | | | | | | | | | | |
| | PET | | | | blue | pink, blue | | | | | | | | | |
| | PETSD | | | | | | | | | | | | | | |

| | |
|---|---|
| *Precipitation* | Annual precipitation |
| | Precip. seasonality |
| | Number of Months with Snow |
| | Aridity Index |

**Supplementary Table 4**: Results from model validation, with highlighted cells showing the 'best' value when comparing the values of each community metric model. The results of mean square error (MSE) following 10-fold cross-validation of (a) the main models, a mixture of sampled soil properties and SoilGrids data and (b) models containing only SoilGrids data. MSE was calculated for the all predicted data ('Total'), then for predicted values the tertiles of the observed data ('Low', 'Mid' and 'High' values). In addition, the $R^2$ squared values of (c) the main models and (d) the models that contained only SoilGrids data. $R^2$ was calculated using the 'MuMIn' package, which calculates the marginal $R^2$ ('Marginal'), i.e., variance explained by the fixed effects, and the conditional $R^2$ ('Conditional'), i.e., variance explained by the fixed and random effects (Barton 2018).

| | Total | Low | Mid | High |
|---|---|---|---|---|
| a) MSE – Main Models | | | | |
| Species Richness | 1.405 | 0.930 | 0.819 | 3.637 |
| Abundance | 18455.07 | 1824.77 | 2505.14 | 50135.34 |
| Biomass | 3224.46 | 250.94 | 382.39 | 8868.71 |
| b) MSE – SoilGrids Models | | | | |
| Species Richness | 1.394 | 0.900 | 0.835 | 3.61 |
| Abundance | 18680.88 | 1660.924 | 2570.47 | 50897.171 |
| Biomass | 3068.31 | 189.37 | 464.13 | 8389.86 |
| | Marginal | | Conditional | |
| c) $R^2$ – Main Models | | | | |
| Species Richness | 0.105 | | 0.741 | |
| Abundance | 0.172 | | 0.611 | |
| Biomass | 0.186 | | 0.612 | |
| d) $R^2$ – SoilGrids Models | | | | |
| Species Richness | 0.112 | | 0.820 | |

| | | |
|---|---|---|
| Abundance | 0.219 | 0.619 |
| Biomass | 0.224 | 0.638 |

### Supplementary Figures

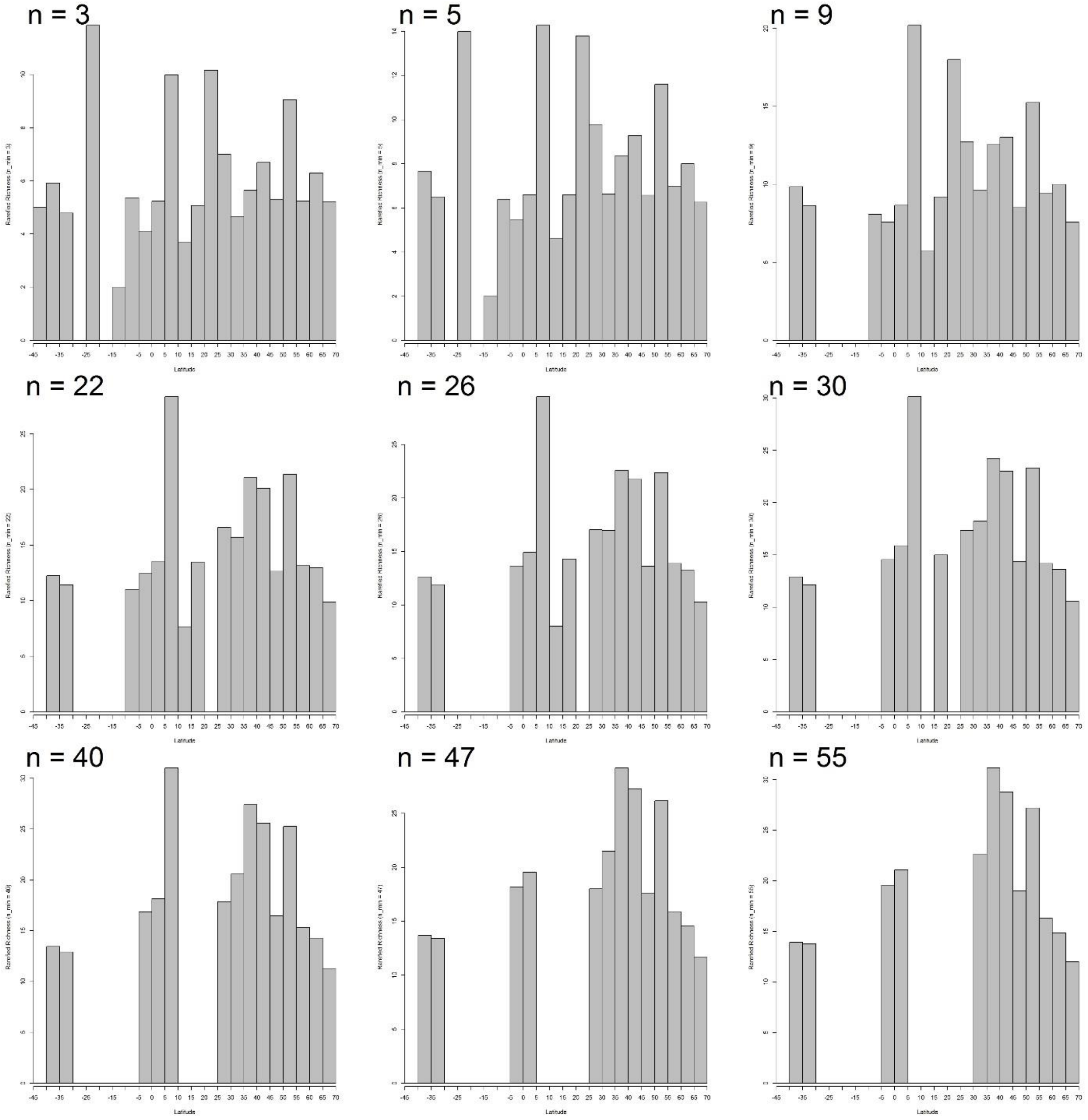


**Supplementary Figure 1**: The changes in the rarefied richness across 5 degree latitude bands, when the number of sites used during the sample-based rarefied richness calculation is changed. Within each band, sample-based rarefied richness was calculated as the mean number of species (including morphospecies, and unique genus', as described in Methods) across 1000 random samples (without replacement) of *n* sites. To test robustness to low number of sampled sites, *n* was varied between 3 and 55).

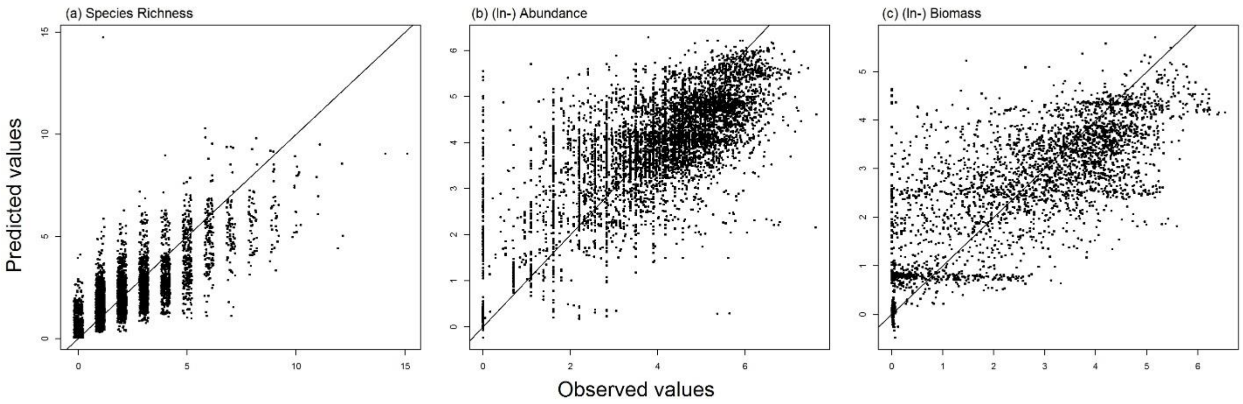


**Supplementary Figure 2:** 10-fold cross validation of the three main community metric models, (a) species richness, (b) abundance and (c) biomass. X-axis shows the observed value, and Y-axis the predicted value, black line is the 1:1 line. The underlying dataset of each model was randomly split into 10 nearly-equal size groups. Using the model structures produced following simplification, the model was built using 9 of the groups of data. The 10th group of data was predicted from the re-built model. This process was repeated until all 10 groups of data had been predicted. The predicted data was plotted against the observed data.

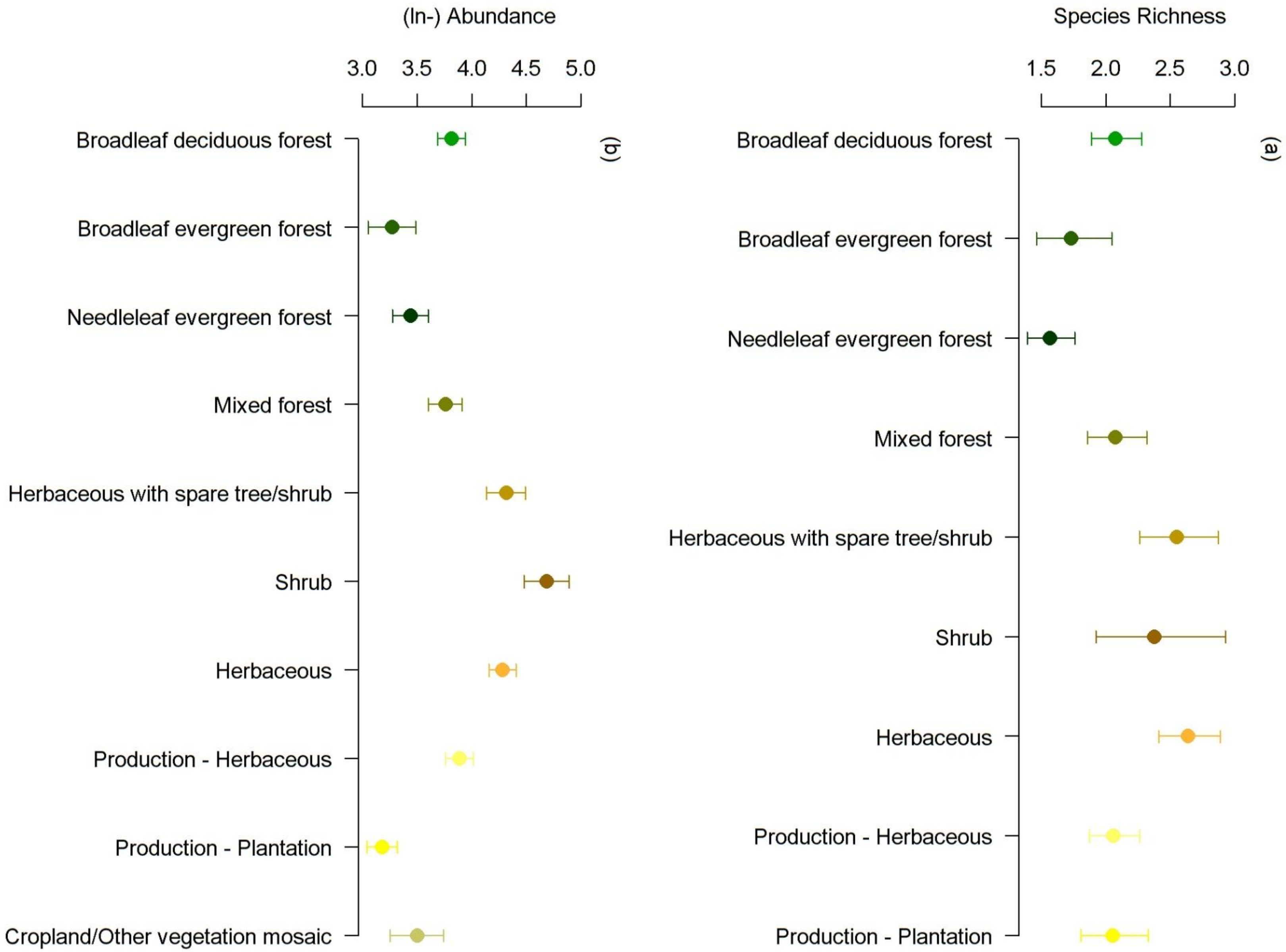
(b)
(ln-) Abundance
3.0
3.5
4.0
4.5
5.0
Broadleaf deciduous forest
Broadleaf evergreen forest
Needleleaf evergreen forest
Mixed forest
Herbaceous with spare tree/shrub
Shrub
Herbaceous
Production - Herbaceous
Production - Plantation
Cropland/Other vegetation mosaic
(a)
Species Richness
1.5
2.0
2.5
3.0
Broadleaf deciduous forest
Broadleaf evergreen forest
Needleleaf evergreen forest
Mixed forest
Herbaceous with spare tree/shrub
Shrub
Herbaceous
Production - Herbaceous
Production - Plantation

**Supplementary Figure 3**: Changes in (a) species richness and (b) ln-abundance across the different habitat cover categories. Values of species richness and abundance are predicted from the main models when all other variables are at zero, i.e., the mean. Error bars show 1 standard error. Not all habitat cover categories had sampled estimates (i.e., species richness could not be estimated for ‘Cropland/Other vegetation mosaic’).

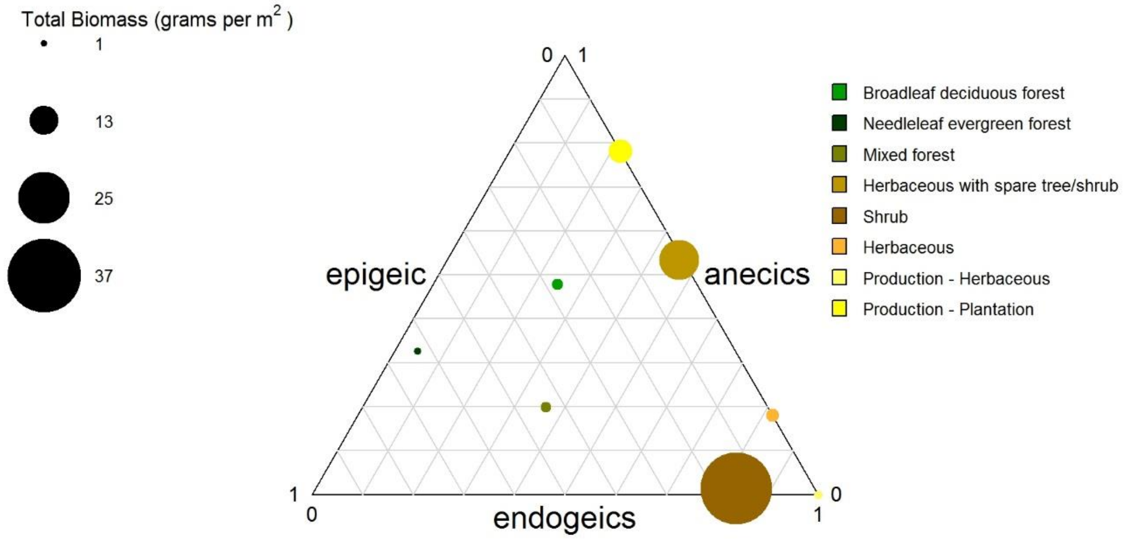


**Supplementary Figure 4**: Composition of the three ecological groups (epigeic, endogeic and anecic earthworms) as predicted by the habitat cover. Circle size is relative to the total biomass predicted for the habitat cover, and circle colour indicates the habitat cover. Position within the three axes indicates the proportion of each of the three ecological groups within the community. Not all habitat cover categories had sampled estimates (i.e., biomass could not be estimated for ‘Broadleaf evergreen forest’ or ‘Cropland/Other vegetation mosaic’).
